# Janus Mazebots and Cellbots Navigating Obstacles in Dense Mammalian Environment

**DOI:** 10.1101/2024.06.05.597538

**Authors:** Max Sokolich, Sudipta Mallick, Calin Belta, Ron Weiss, Sambeeta Das

## Abstract

The field of microrobotics has immensely grown in the last few decades, exhibiting several challenges as new features such as shapes, sizes, and actuation mechanisms are explored. Two of the biggest challenges faced in microrobotics are the development of a control system suited for precise microrobotic manipulation, and the ability to navigate microrobots in densely populated environments. In this paper, we fabricate the Mazebot microrobots using silica spheres and ferromagnetic coating, and we use them to create cellbots with genetically modified Chinese Hamster Ovary (CHO) cells. Subsequently, we navigate both the Mazebots and the cellbots through a dense environment populated by CHO cells. The Mazebots navigation is done with a control system that allows the Mazebots to swim on their own, or guide a specific cell from a given origin to a target location while avoiding cell obstacles. The control system operates in open and closed-loop modes, where the first one allows the microrobot to reorient the cell using self-induced fluid vortices, and the second one closely follows a predefined trajectory along the origin and destination. On the other hand, the cellbots navigation is done in closed-loop operation. This enables cell manipulation for potential applications in cell and tissue engineering when in a confined space. Biocompatibility of the Mazebots is confirmed through the exposure of CHO cells to the robots for 24 hours. Experimental results demonstrate the functionality of our algorithm and its potential for biomedical applications, showcasing our system as a powerful and efficient solution for precise cellular manipulation.

## INTRODUCTION

The field of micro-robotics has expanded significantly as robots, electronic devices, and actuation systems have steadily decreased in size.^1–4^ This growth has fostered a diverse interdisciplinary research area that merges robotics, automated control, material science, chemistry, biology, and biomedicine. The inevitable miniaturization of robotics holds great promise for various applications such as environmental monitoring, biomedical interventions, and manufacturing sensing.^5–9^

As researchers and engineers continue to explore and innovate in microrobotics, several inherent challenges regarding actuation and control have been addressed. Hence, functional structures powered by external stimuli have been proposed as solutions, including magnetic field, light, chemical fuel, electric field, and acoustic waves,^10–18^ where multiple stimuli can be simultaneously employed to improve navigation of microrobots. ^10,19,20^ In healthcare, tissue engineering, and biomedical applications, the preference is for energy sources that enable remote control while harmlessly penetrating complex structures. Thus, magnetic microrobots have gathered attention due to their biocompatibility, responsiveness, and ease of operation.^6,12,13,21–23^ Then, magnetically actuated microrobots are suitable for several biomedical applications such as drug delivery,^24^ cargo-transport,^25^ and single-cell manipulation,^10,26,27^ where the latest has historically been of great interest for biologists. Single-cell manipulation is crucial for understanding cellular-level heterogeneity within complex structures,^28^ which can lead to accurate diagnosis and therapeutic advancements.

Despite extensive research, achieving reliable single-cell manipulation remains a complex and overarching challenge. Techniques like pipettes,^29^ optical tweezers,^30^ and microfluidic systems^31^ have been explored for single-cell manipulation. Microfluidic methods have gained attention for cell isolation, patterning, transportation, and some other applications,^20,32–34^ but suffer from high costs, specific flow rate requirements, and challenges in cell culture and fluid manipulation. ^31^ In response, magnetically actuated microrobots are emerging as an option to overcome single-cell manipulation challenges.^11,35**?**^ While cellular manipulation is one of the most promising applications within microrobotics,^36^ achieving precise and independent single-cell manipulation remains an open challenge since this requires microrobots to be sufficiently small to interact one-on-one with single cells, and reliable enough to function in dense cellular environments. Current magnetic microrobots for cell manipulation exceed 100 *µ*m in size,^37–39^ primarily operating in obstacle-free settings due to the complexities of actuation for smaller microrobots in crowded environments. Moreover, the transportation of genetically modified cells, integral to genetic circuits for cell engineering,^40,41^ presents another opportunity for microrobot application. Recent developments in cellbots or immunobots^42–44^ demonstrate promise, where biocompatible magnetic microrobots are encapsulated within cells, directing cellular motion.

This study presents the fabrication, control, and actuation of “Mazebots”, spherical microrobots of 4.8 *µ*m diameter. Mazebots are fabricated from silica microspheres, embedded with magnetic properties for responsiveness and controllability. We also engineered Cellbots integrating genetically modified sender Chinese Hamster Ovary (CHO) cells with Mazebots. Furthermore, we present a versatile control system capable of both open and closed-loop operation modes, tailored to specific applications. Closed-loop control is explored for independent Mazebot navigation and Cellbot manipulation, while open-loop control facilitates single-cell manipulation by Mazebots. All applications are tested in dense cellular mammalian environments, showcasing the potential of the results for both, *in vivo* and *in vitro* implementations.

## Results and Discussion

Figure 1a shows the experimental setup. A 3D electromagnetic manipulation Helmholtz coil system was designed to output uniform magnetic fields in 3 Dimensions. The coils are mounted on a Zeiss Axiovert microscope. Figure 1b shows the a visual illustration of the workspace and an overview of the closed loop control methodology.

**Figure 1.**
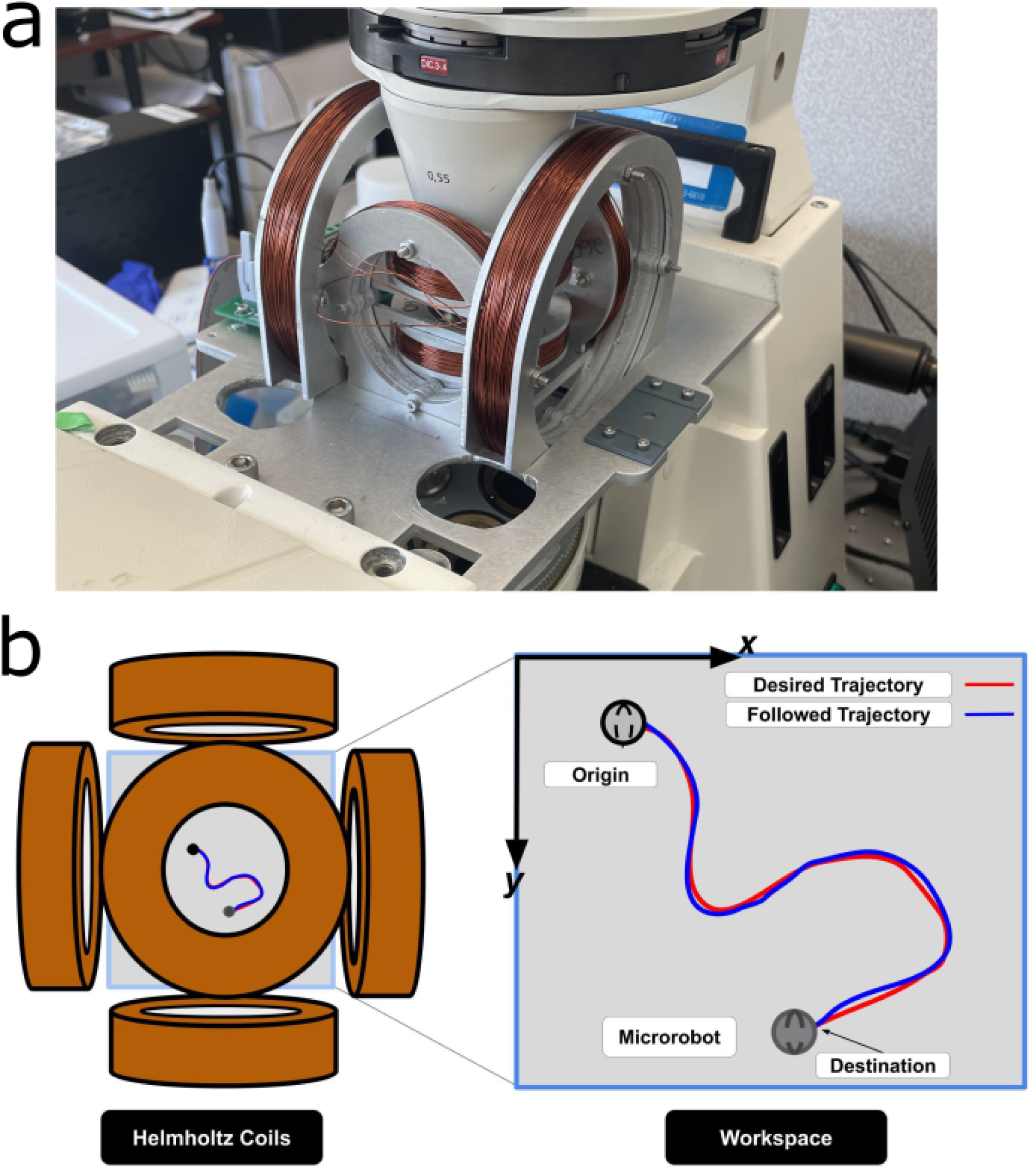
Experimental schematic illustration demonstrating the closed loop control of the microrobots using a 3 dimensional Helmholtz system.

Figure 2a shows the block diagram for autonomously controlling the microrobots and cellbots. First, images are captured from a microscope camera and read using OpenCV python. Next, a custom written tracking and detection algorithm determines the selected microrobots position and the surrounding cellular obstacles. Next, a path or trajectory is either manually determined through observation or through the RRT path planning algorithm. Finally, the necessary magnetic field action commands are applied to the electromagnetic coil system to guide the microrobot along the desired path.

**Figure 2.**
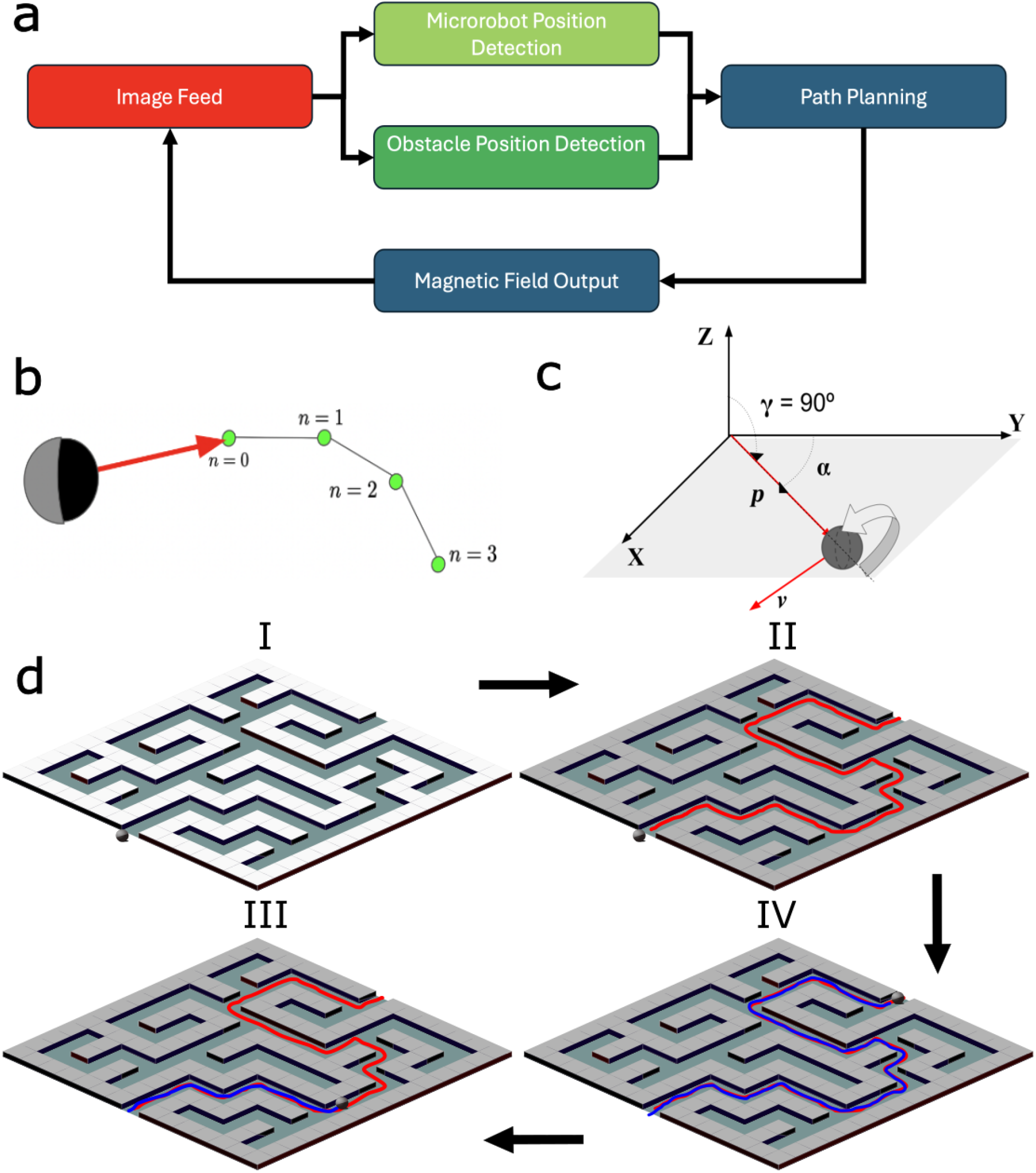
Schematic of Mazebot rolling on a glass surface (gray plane) actuated by a rotating magnetic field, where **p** represents the position of the microrobot, **v** the velocity, *α* the polar angle and *γ* the azimuth angle, which determines the rolling axis.

Figure 2b demonstrates the procedure for guiding the microrobot or cell-bot along the calculated trajectory. Each trajectory consists of N nodes in the form of Cartesian coordinate pixel locations T = [(x1, y1), (x2, y2), …, (xN, yN)]. The error between the microrobots current position (xr, yr) and the next node (n = 0) is calculated. Furthermore, the direction vector (in red) pointing from the microrobots current position (xr, yr) to the next node (n = 0) is also calculated. The algorithm described in more detail in a later section then aims to reduce the error between the microrobots current position and the nodes in the trajectory array.

Figure 2c describes the microrobot or cellbots actuation system. The microrobot and cellbots described in this paper are actuated via a rotating magnetic field. Equations 1, 2, and 3 generate a rotating magnetic field in time. There are 3 input parameters that control the motion of the microrobot or cellbot. 0 *< α <* 360 is the polar angle or steering angle of the microrobot that allows it to travel in any direction along the 2D plane. 0 *< γ <* 180 is the azimuthal angle and determines the rotation axis in 3 dimensions. *γ* = 90 throughout this paper to keep the microrobot or cell bot rolling along the xy plane. Finally, 0 *< ω <* 20 defines the rotating magnetic field frequency that defines the robots speed. Speed characteristic studies can be found in a later section.

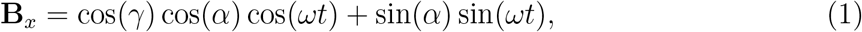

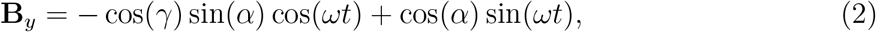

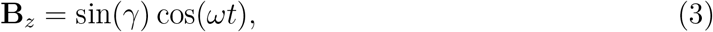

### Micro-robot Control

Two methods for automated control of microrobots and cellbots are now described. The first is manual trajectory drawing and the second is automated trajectory calculating. For the manual trajectory drawing, the user can draw a path with the right mouse button that avoids the cells through observation. For automated trajectory calculation we used the RRT path planning algorithm to generate automatic trajectories that avoid nearby cells. Rapidly Exploring Random Trees (RRT) is a popular motion planning algorithm. It is used to efficiently explore and plan paths for a robot or other agents in a continuous high dimensional space. The primary idea behind RRT is to incrementally build a tree structure through random sampling and exploration of the configuration space. First a black and white mask is created to clearly create contrast with the nearby cells. The effectiveness of the mask can be controlled by adjusting three parameters; mask threshold, mask blur, and mask dilation. The mask is generated by first converting the color image to gray-scale. Then, the user can blur the gray-scale image using the opencv function cv2.blur. This helps reduce noise, eliminate small details, and create a more simplified version of the image. Then, a threshold is applied using cv2.threshold function. The function is used to apply a fixed-level threshold to each pixel in an image which converts the image into black and white. This results in a black and white mask with the cells in white and the traversable region in black. Finally, the user can apply cv2.dilate to the mask to dilate the cellular white regions. This creates a safety zone surrounding the cells that prevents the microrobots from getting too close to the cells.

#### Algorithm 1

RRT Path Planner^17^

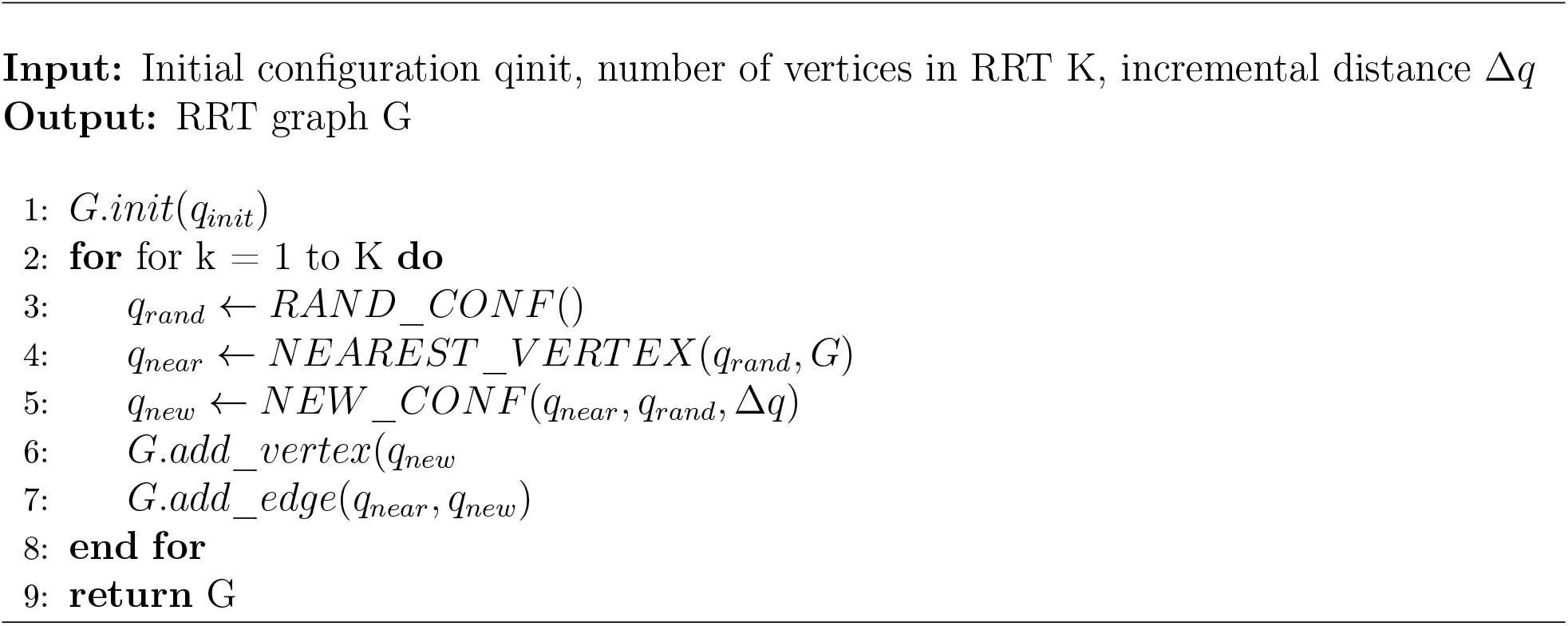

Once a mask is calculated, the user can click on a desired microrobot, then click on a target location within the black traversable region, and the RRT algorithm will automatically generate a path that avoids the white cellular regions. A visual schematic of the procedure is outlined in figure 3.

**Figure 3.**
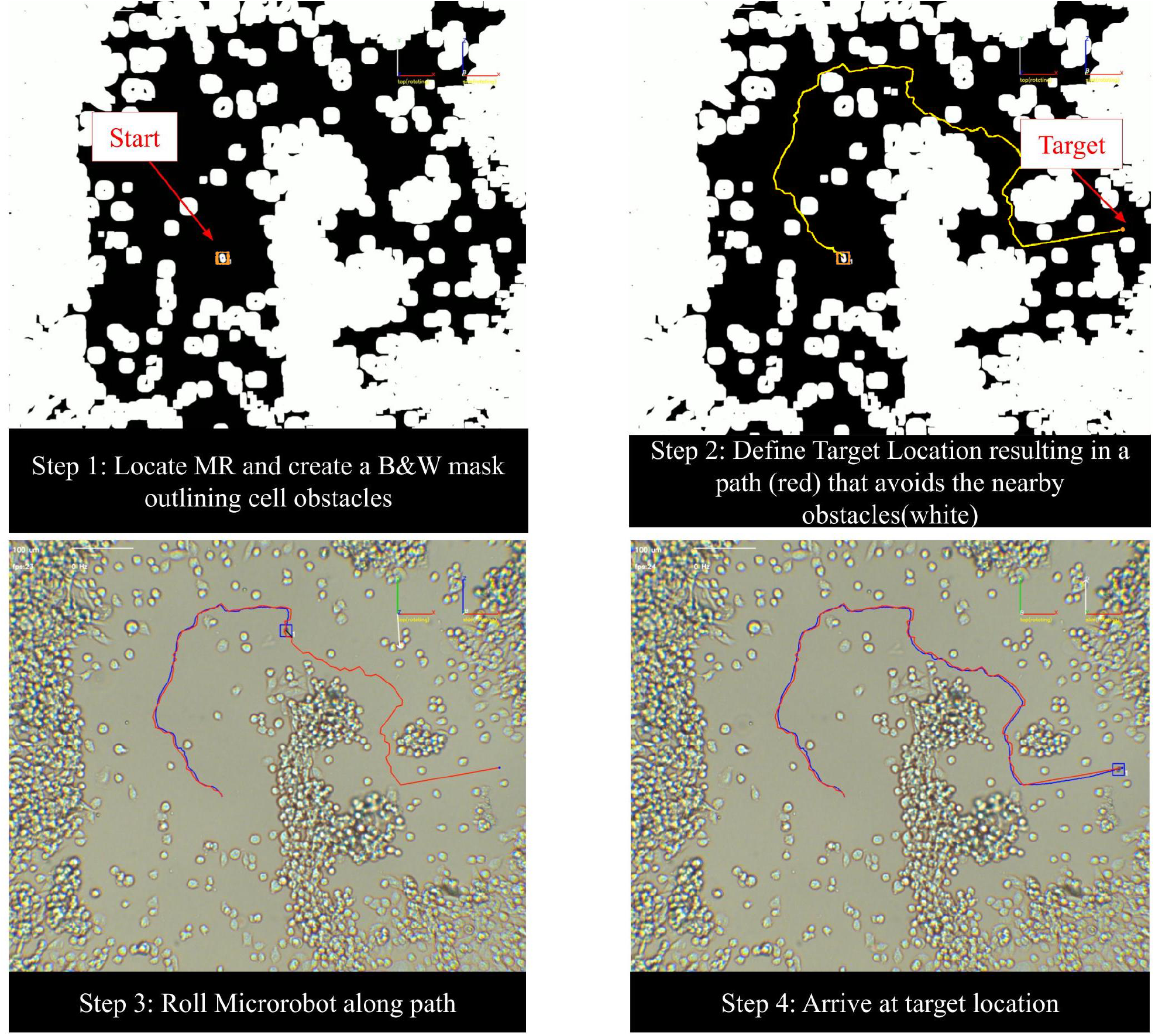
Path planning steps in an experiment. Step 1 consists of locating the microrobot and creating a mask that outlines the cells in white. Step 2 consists of defining the target location and RRT generated path or manually drawing the path with the mouse. Step 3 and 4 then result in rolling the microrobot along the defined trajectory and arriving at the target position.

Algorithm 1 describes the closed loop control procedure for guiding the microrobot or cellbot along the calculated path described above. First, the trajectory node array is defined with nodes equal to the Cartesian pixel coordinates of the path. Next, a threshold variable δ is defined that describes a distance in pixels. Next, for each incoming image frame, we calculate the error between the robots current position [*p*_*x*_, *p*_*y*_] and the current node [*x*_*i*_, *y*_*i*_].

This is essentially the distance 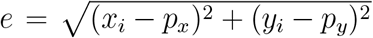 between the two. Next, *α* or the steering angle is calculated by the angle made by the vector extending from the robots current position to the current node. We then output a rotating magnetic field defined by equations 1,2, and 3 above using the calculated *α*. If the error *e* is less than the defined threshold δ, then we say the robot has arrived at the current node, and to move on to the next node. Then the procedure repeats until all nodes in the array have been explored.

#### Algorithm 2

Nonlinear closed-loop trajectory tracking for rolling microrobots

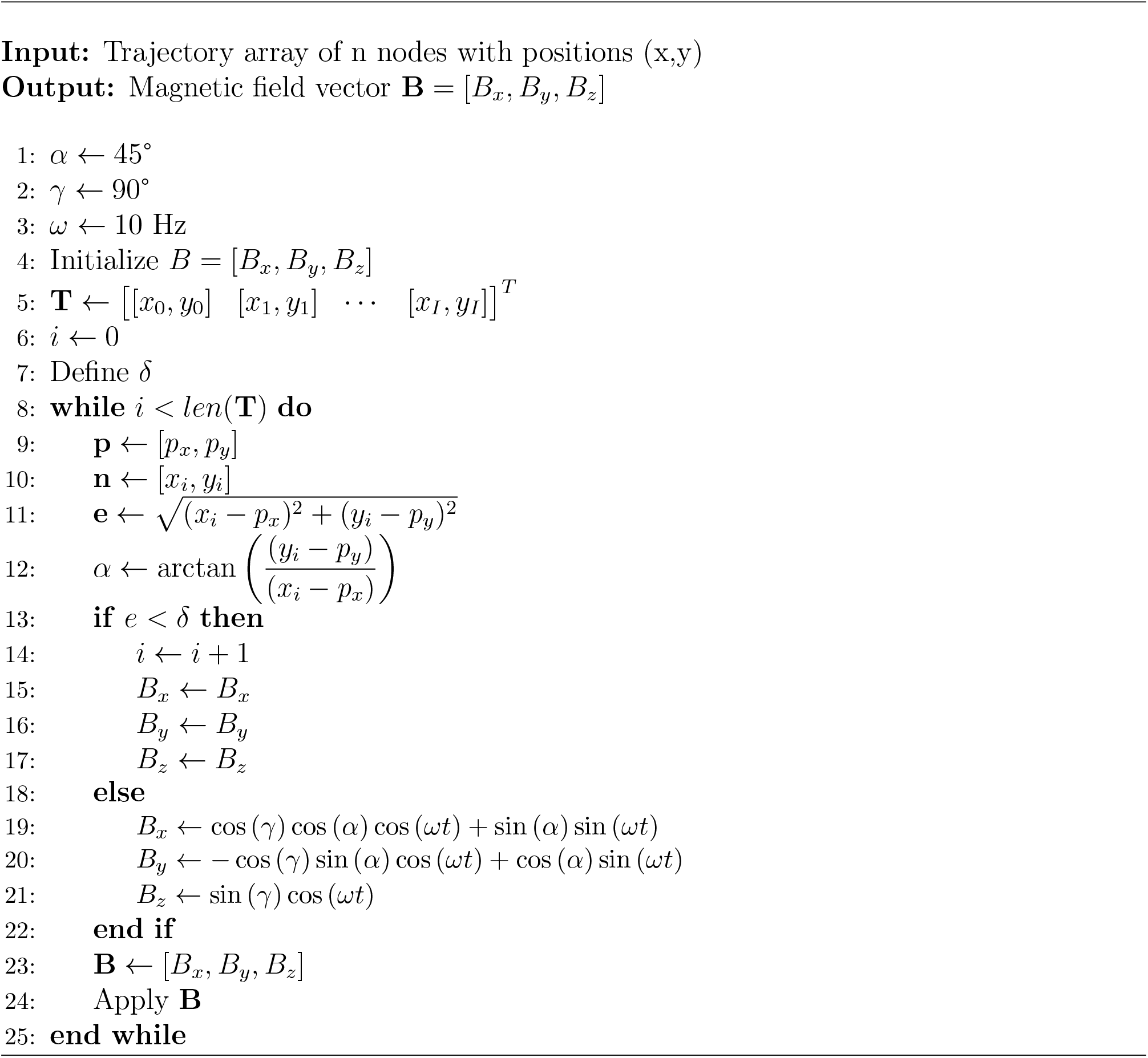

## Discussion

Figure 4 illustrates the algorithms implementation in experiment. An artificial maze was designed and fabricated using standard photo lithographic mask alignment protocols using SU8 photo-resist. In figure 4a, a cellbot containing three 5um silica microrobots coated in 100nm Nickel can be seen autonomously solving the complex maze. The cellbot (approximately 15 um in size) begins at the top left corner and solves the maze in approximately 108 seconds at 10 Hz and 5 mT. Similarly, in Figure 4b, a 20um Silica microrobot coated in 100nm Nickel can be seen autonomously solving the maze in approximately 56 seconds at 10 Hz and 5 mT.

**Figure 4.**
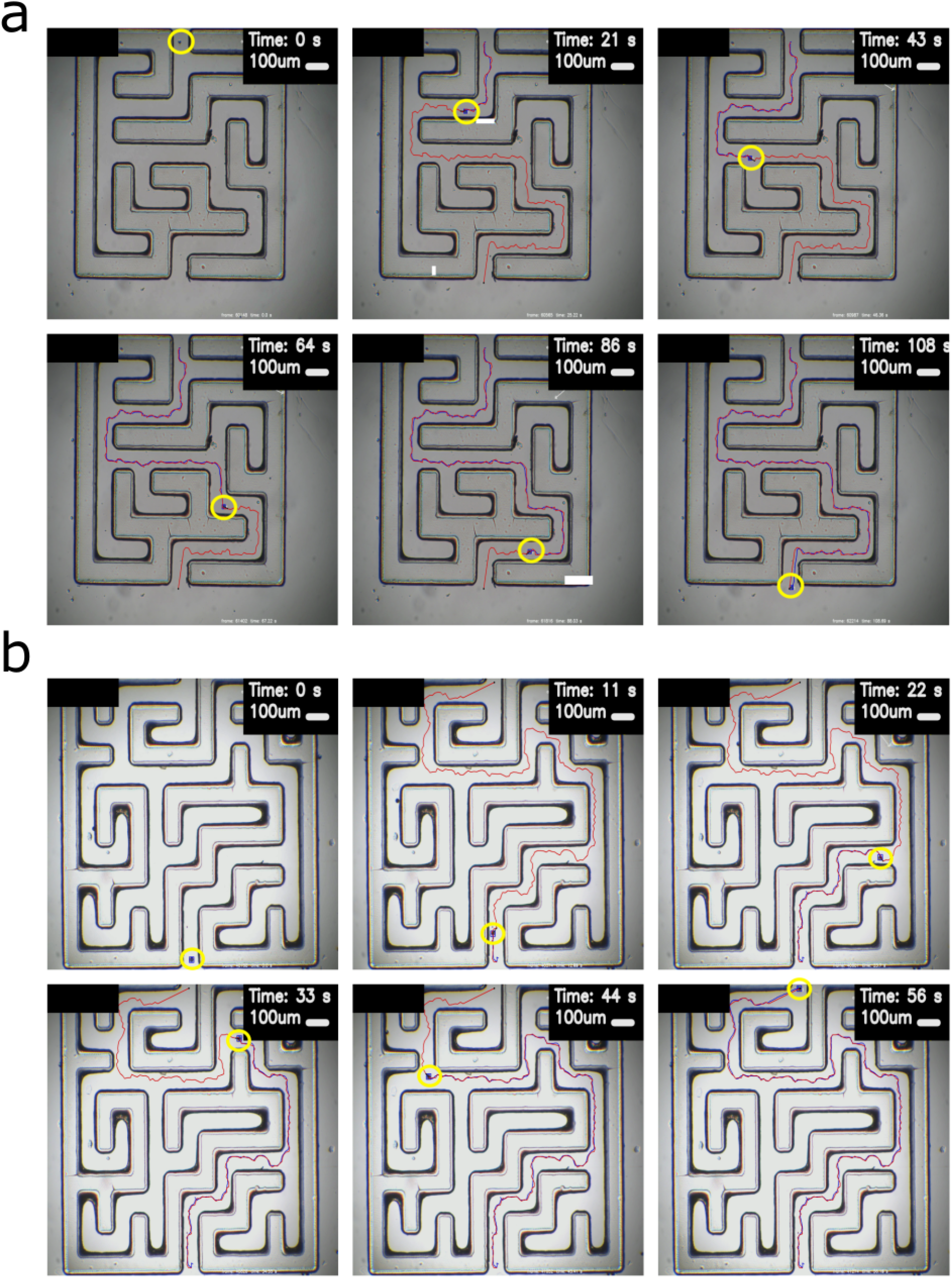
a) 3 bot Cellbot autonomously solving the maze. b) 20um silica microrobot autonomously solving the maze.

Additionally, experiments were conducted in a dense cell environment. Figure 5a shows a 5um silica microrobot coated in 100nm nickel navigating the cellular environment along a manually drawn trajectory (red). Figure 5b shows a cell bot with four 5 um silica microrobots autonomously avoiding cells via the RRT algorithm. Furthermore, it was found that both cellbots and microrobots were able to manipulate single cells under a rotating magnetic field. A gaming joystick is used to manually manipulate single cells by pushing the cells.

**Figure 5.**
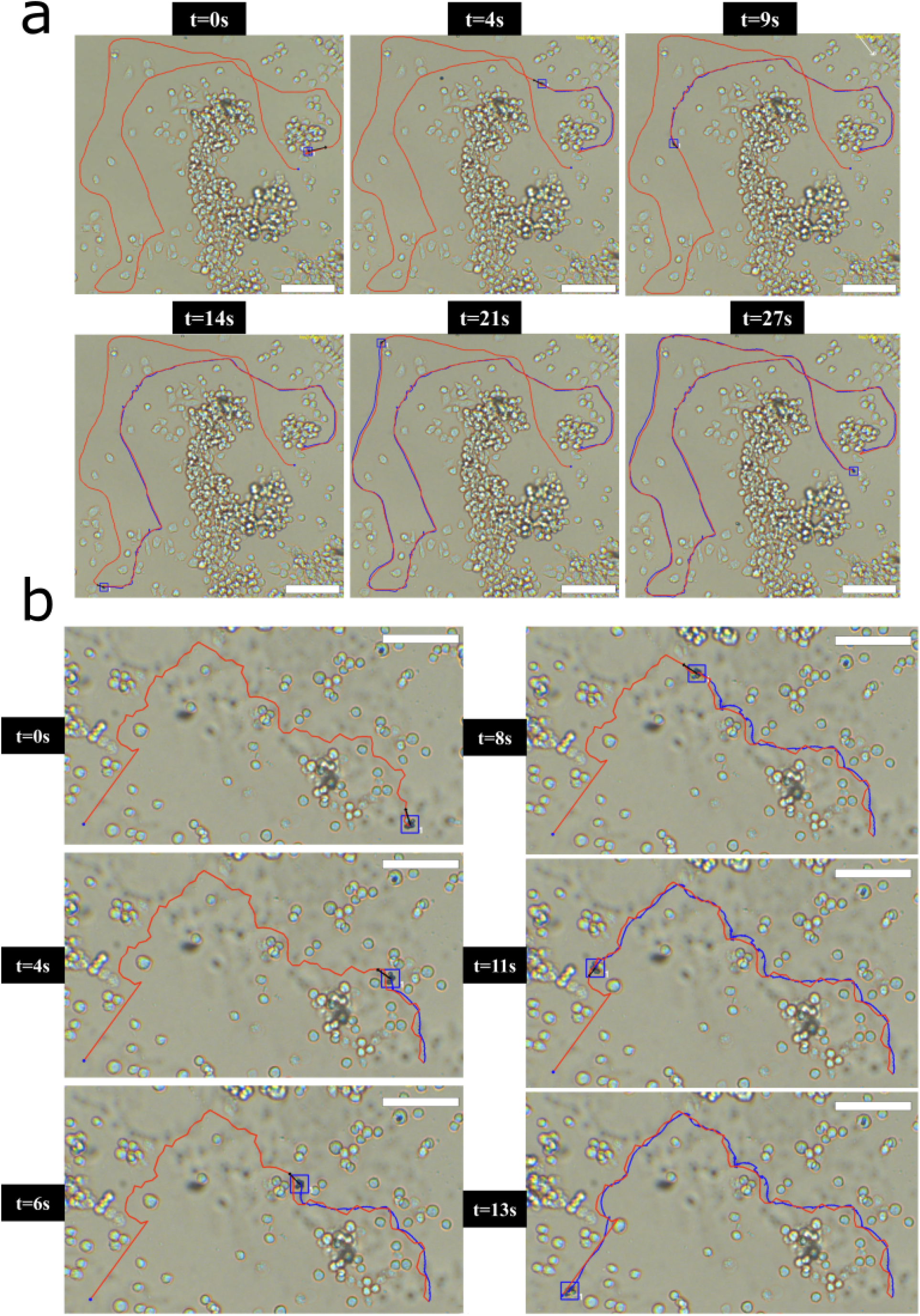
a) 5um Janus rolling microrobot navigating around cells along manually drawn trajectory. Scale bar = 100 um. b) Cellbot autonomously navigating around cells using RRT generated path.

**Figure 6.**
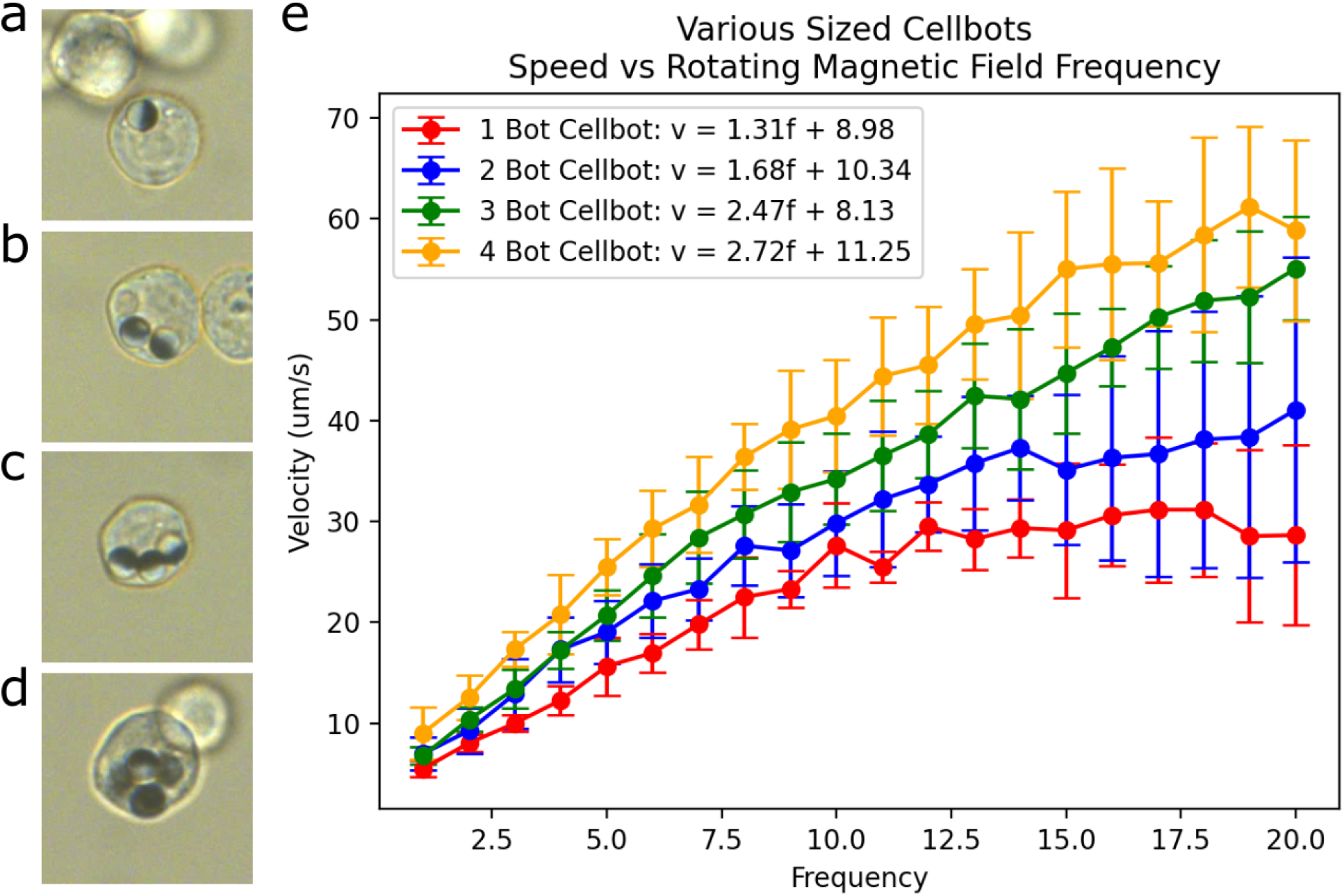
Images of cell bots with various number of 5um silica Janus microrobots inside. a) 1 robot Cellbot. b) 2 robot Cellbot. c) 3 robot Cellbot. d) 4 robot Cellbot. e) Cellbots speed (um/s) vs rotating magnetic field frequency (Hz) at 5mT.

### Single Cell Manipulation

It was found that both Janus rolling microrobots and cellbots were capable of manipulating single cells around the workspace. By

### Speed Characteristic

Figures 7 and 8 describe the relationship between the speed of the microrobot and the rotating magnetic field frequency. The magnetic field intensity was set at 5 mT for the speed Characterization experiments. Figure 7a shows a microscope image of a 5um silica microsphere coated in 100nm Ni. Figure 7b shows a microscope image of a 20um silica microsphere coated in 100nm Ni. Figure 7c shows a microscope image of a 5um silica dimer microsphere coated in 100nm Ni. Figure 7d shows a microscope image of a 20um silica Dimer microsphere coated in 100nm Ni. It is clear from the graph that Dimer microrobots, that is, microspheres attached to one another move significantly faster than a single microrobot. It was found that the 20 um silica Dimer microrobot moved at the highest speeds reaching almost 400 um/s at 20 Hz. A linear regression was performed on the data and a best fit line was calculated to be *v* = 1.04*f* + 3.18 where v is the robots velocity and f is the rotating magnetic field frequency. Next the 20 um silica Dimer microrobot was the next fastest reaching speeds close to 200 um/s at 20 Hz. The equation describing this microrobots motion is *v* = 8.17*f* + 17.27. Next the 5um Dimer microrobot demonstrated the 3rd fastest microrobot of the group reaching a top speed of approximately 70 um/s. The speed vs frequency relationship for this microrobot is *v* = 3.12*f* + 5.07. Finally, the single 5um silica microrobot was the slowest of the 4 with a top speed of approximately 24 um/s and a best fit line characterized by *v* = 1.04*f* + 3.18. From the data, it is clear that larger particles move faster than smaller particles at the same rotating magnetic field frequency and magnetic field strength. Furthermore, microrobot Dimers or microrobots attached to one anther to form a doublet microrobot also move faster than their single counterparts.

Figure 8 shows a similar speed vs frequency characterization study for cell bots containing a different numbers of 5um microrobots inside. Figure 8a shows a microscope image of a cellbot containing 1 microrobot inside. Figure 8b shows a microscope image of a cellbot containing 2 microrobots inside. Figure 8c shows a microscope image of a cellbot containing 3 microrobots inside. Figure 8d shows a microscope image of a cellbot containing 4 microrobots inside. Figure 8e displays the relationship between cellbot speed, the rotating magnetic field frequency and the number of microrobots ingested. Linear regressions were also performed on the Cellbot data. Cellbots with 4 microrobots ingested move the fastest. The speed vs frequency relationship for 4 bot cellbots is *v* = 2.72*f* + 11.25. The next fastest cellbot was the cellbot with 3 microrobots ingested. The best fit line is characterized by *v* = 2.47*f* + 8.13. Next, cellbots with 2 microrobots ingested moved the 3rd fastest with a speed vs frequency relationship described by *v* = 1.68*f* + 10.34. Finally, cellbots with a single microrobot ingested unsurprisingly moved the slowest with a best fit line described by *v* = 1.31*f* + 8.98. It can be seen by figure 8e that the speed of cellbots is proportional to the rotating magnetic field frequency. Furthermore, the speed of the cell-bots also depend on the number of microrobots ingested, where more ingested microrobots result in faster cellbots. However, it is observed that a cell can die if too many microrobots are ingested. As a result, future work aims to understand the relationship between the number of cellbots ingested and the the viabilty of the cell over time.

## Conclusion

The Mazebots fabricated in this work exhibited the desired magnetic properties, successfully responding to the proposed actuation mechanism. Furthermore, the Mazebot system showed compatibility with the cells, as confirmed through cell toxicology experiments, establishing its bio-compatibility and suitability for biological and biomedical applications, such as the fabrication and manipulation of the Mazebot-driven Cellbots created in this study.

Moreover, the proposed control system can operate in both, closed and open-loop modes, fulfilling needs for different applications, such as the sole Mazebot automatically navigating the maze environment, the transportation and reorientation of the cell using self induced vortexes or the autonomous movement of the Cellbots. In addition, our control algorithm offers advantages over traditional PID implementation. Its inherent adaptability and self-adjusting features, allow the Mazebot to navigate its path with higher levels of accuracy, without the complexities associated with PID tuning. This reduces the potential for human error and makes the Mazebot system suitable and reliable for several future applications. It is important to note that the path generated is not an optimal path that minimizes the number of cellular collisions nor time taken to complete the path.

## Methods/Experimental

### Synthesis of Microrobots

5um Silica microspheres were purchased from Bang Laboratories Inc (Catalog Number SSD5003) in a dry powder form. 20um Non-functionalized Silica microspheres where purchased from Alpha Nanotech (Batch Number #42222) suspended in Milli-Q water. Next, microspheres were drop casted on a clean glass slide and allowed to dry. Next, the glass slide was place in a dual electron beam vapor deposition chamber (PVD Products) and 100 nm of Nickel was deposited on the surface. Janus microrobots were then scratched from the surface and injected into the experimental environment.

### Synthesis of Cellbots

Cellbots were fabricated using 5 um Silica Ni coated microrobots described above. They were scratched from the surface and introduced to a culture of CHO cells. After approximately 24 hours, cells ingest the microrobots and become cellbots.

### Electromagnetic Manipulation System

The electromagnetic manipulation system consists of a desktop computer (OMEN by HP Obelisk Desktop PC), a Micro-controller (Arduino Mega 2560), 6 motor drivers (BTS7960 43A H-Bridge), a Benchtop power supply, CCD camera (BFS-U3-50S5C-C USB 3.1 Black-fly® S, Color Camera) and a 3 dimensional Helmholtz setup. Custom control software was written in Python that received live image feed from the camera and transmitted magnetic control signals to the Arduino and coil system. A rotating magnetic field was applied resulting in a uniform magnetic field of 5 mT.

### Fabrication of Artificial Maze

The first step for maze fabrication is cleaning the substrate, which is the glass slide, in the solution of acetone (Product No.: 2462) and IPA (Product No.: 9088) with sonication accordingly. Bake the substrate at 200°C for 5 minutes to improve the photoresist adhesion after cleaning. The next step is to coat the substrate with photoresist SU8-2050. 1mL of SU8-2050 is carefully poured at the center of the substrate to prevent any glass bubbles generated. The glass slide is then spun at 500 rpm for 60 s followed by 1500 rpm for 60 s to acquire a thickness at around 120um. To have a smooth surface, the substrate is then placed at a horizontal surface for 24h after spinning. The pre-exposure bake is heating the substrate at 65°C for 5 min and 95°C for 20 mins. Using the mask aligner (NXQ8006) to expose the substrate with a dose of 240 mJ/cm2 to make the place where the wall of the maze cross-linked. Post-exposure bake is baking the sample at 65°C for 5 min and 95°C for 10 mins. The sample was then developed in SU8 developer (CAS No.: 108-65-6) for 10 min to wash out the photoresist that is not cross-linked with the glass slide.

